# Constructing and Visualizing Cancer Genomic Maps in 3D Spatial Context by Phenotype-based High-throughput Laser-aided Isolation and Sequencing (PHLI-seq)

**DOI:** 10.1101/278010

**Authors:** Sungsik Kim, Amos Chungwon Lee, Han-Byoel Lee, Jinhyun Kim, Yushin Jung, Han Suk Ryu, Yongju Lee, Sangwook Bae, Minju Lee, Kyungmin Lee, Ryong Nam Kim, Woong-Yang Park, Wonshik Han, Sunghoon Kwon

## Abstract

A spatially resolved analysis of the heterogeneous cancer genome, in which the data are connected to the three-dimensional space of a tumour, is crucial to understand cancer biology and the clinical impact of cancer heterogeneity on patients. However, despite recent progress in spatially resolved transcriptomics, spatial mapping of genomic data in a high-throughput and high-resolution manner has been challenging due to current technical limitations. Here, we describe a novel approach, phenotype-based high-throughput laser-aided isolation and sequencing (PHLI-seq), which enables high-throughput isolation of a single-cell or a small number of cells and their genome-wide sequence analysis to construct genomic maps within cancer tissue in relation to the phenotypes of the cells. By applying PHLI-seq, we reveal the heterogeneity of breast cancer tissues at a high resolution and map the genomic landscape of the cells to their corresponding spatial locations and phenotypes in the tumour mass. Additionally, with different staining modalities, the genotypes of the cells can be connected to corresponding phenotypic information of the tissue. Together with the spatially resolved genomic analysis, we can infer the histories of heterogeneous cancer cells in two or three dimensions, providing significant insight into cancer biology and precision medicine.

---

Recent advances in sequencing technology have revolutionized oncology and provided an opportunity to overcome the limitations of pathological analysis^1^. However, the mutagenesis and genetic evolution in the tumour mass^2^, resulting in intra-tumour heterogeneity, impede precise deciphering of a causation underlying carcinogenesis. Deep sequencing and computational analysis were proposed to analyse the subclonal population in a tumour mass, each subclone of which could be generated through subclonal expansion of a progenitor tumour cell that acquired advantageous fitness after undergoing an oncogenic initial mutation^3–5^. Instead of sequencing separately the genomic DNA of cells from different subclones in a tumour mass, those approaches relied upon computational inference to separate subclones from the mixed population. Therefore, they cannot elucidate the direct relationship between the genomic data and tissue context during the process. Moreover, because the heterogeneous subclones were sequenced in a pool, the detection sensitivity of variants with a low-level allele fraction was low. To overcome these limitations, other researchers adopted multi-region sequencing or laser capture microdissection instead of sequencing the whole tumour^6–8^. However, these procedures are low throughput and are usually used to process a large number of cells at once (>1000). The ultraviolet (UV) beam used in laser capture microdissection not only hinders the throughput by burning the periphery, but also causes damage to cells^9^. Even in a microdissection using infrared (IR), which uses direct contact methods, cell debris or non-selected cells may cross-contaminate the targeted cells^10,11^. These problems limit the use of microdissection for spatial mapping of the heterogeneous cancer genome in a high-throughput and high-resolution manner. More recently, to provide a higher throughput and sensitivity, a single-cell analysis was proposed as a solution to thoroughly characterize the subclones and variants with a low-level allele fraction in tumours^12–15^. For example, a model of tumorigenesis and the evolutionary history of subclones in a tumour mass could be inferred by highly multiplexed single-cell genome analysis^16^.

Despite the remarkable advances in sequencing technology, the foundations of modern oncology lie in histopathology, which remains the gold standard for comprehending tumorigenesis, relapse, metastasis, cancer evolution, and appropriate clinical applications^17^. Pathological assays, such as histological staining, immunohistochemistry (IHC) staining, and fluorescence in situ hybridization (FISH), can directly provide the molecular information for single cells and their microenvironment. However, the conventional histopathological analyses often fail to identify subclones of cancer or rare cell types, and the subjective interpretation of the histopathological findings may lead to misunderstandings of cancer. For a more precise understanding of cancer and to improve diagnostic performance, it is important to link the spatial and phenotypic histopathological information to the objective and massively parallel measurement of the genomic alteration status of cells in a high-throughput manner^18^. However, the huge amounts of sequencing data cannot be connected to the spatial and phenotypic histopathological information because many cells must be pooled or dissociated in solution before sequencing. This loss of information for linking the subclonal genomic heterogeneity with the histopathological context hinders deeper analyses of the tumour mass. Spatially resolved genomics has, therefore, emerged to address this issue. Considering recent progress in spatially resolved transcriptomics^19–22^, technical advances are required to map the genomic data for cancer spatially.

Here, we describe phenotype-based high-throughput laser-aided isolation and sequencing (PHLI-seq), which can efficiently detect tumor subclonality and variants with low-level allele fractions with high sensitivity and accuracy while preserving information about the 3D spatial organization and histopathological phenotypes of cells by multi-staining modalities. Fully utilizing the spatial and phenotypic information^23,24^, we computationally or pathologically grouped cancer cells in tumour tissue sections into potential subclones. With the group information for the cells as guidance, cell clusters consisting of 20 to 30 neighbouring cells were selected to cover all the groups and effectively represent the spatial organization of the tumour. Next, each cell cluster was physically isolated into different tubes by discharging the region of each cell cluster on an indium tin oxide (ITO)-coated glass slide using an infrared laser pulse. Finally, the extracted DNA from each tube was amplified and sequenced. The procedures, including grouping, selecting, and isolating cells, were automated with custom software to achieve a high-throughput analysis. By establishing this novel approach, we identified distinct subclones and their somatic variants in breast cancer tissues. Overall, we were able to map the heterogeneous genomic landscape of the subclones directly to the spatial and phenotypic organization of the tumour mass. We constructed genomic maps in spatial context in two- or three-dimensional space of cancer tissues, and augmented the dimension of the histopathology and genomics to understand the evolutionary histories of cancer.

## RESULTS

### Workflow of Phenotype-based High-throughput Laser-aided Isolation and Sequencing

For PHLI-seq, tissue sections or cells were stained on a transparent discharging layer-coated glass slide (**Fig. 1a** and **b**). To group the cells in the sample, we segmented the image of the tissue into cell clusters using a conditional random field algorithm. Then, several features, including locational information for the histopathological section and the morphological information for the cell cluster, were extracted from each of the segmented cell clusters. Finally, we generated groups based on the location and morphology of the cell clusters (**Fig. 1a** and Online Methods), and the cell clusters to be sequenced were chosen to represent all groups.

**Figure 1.**
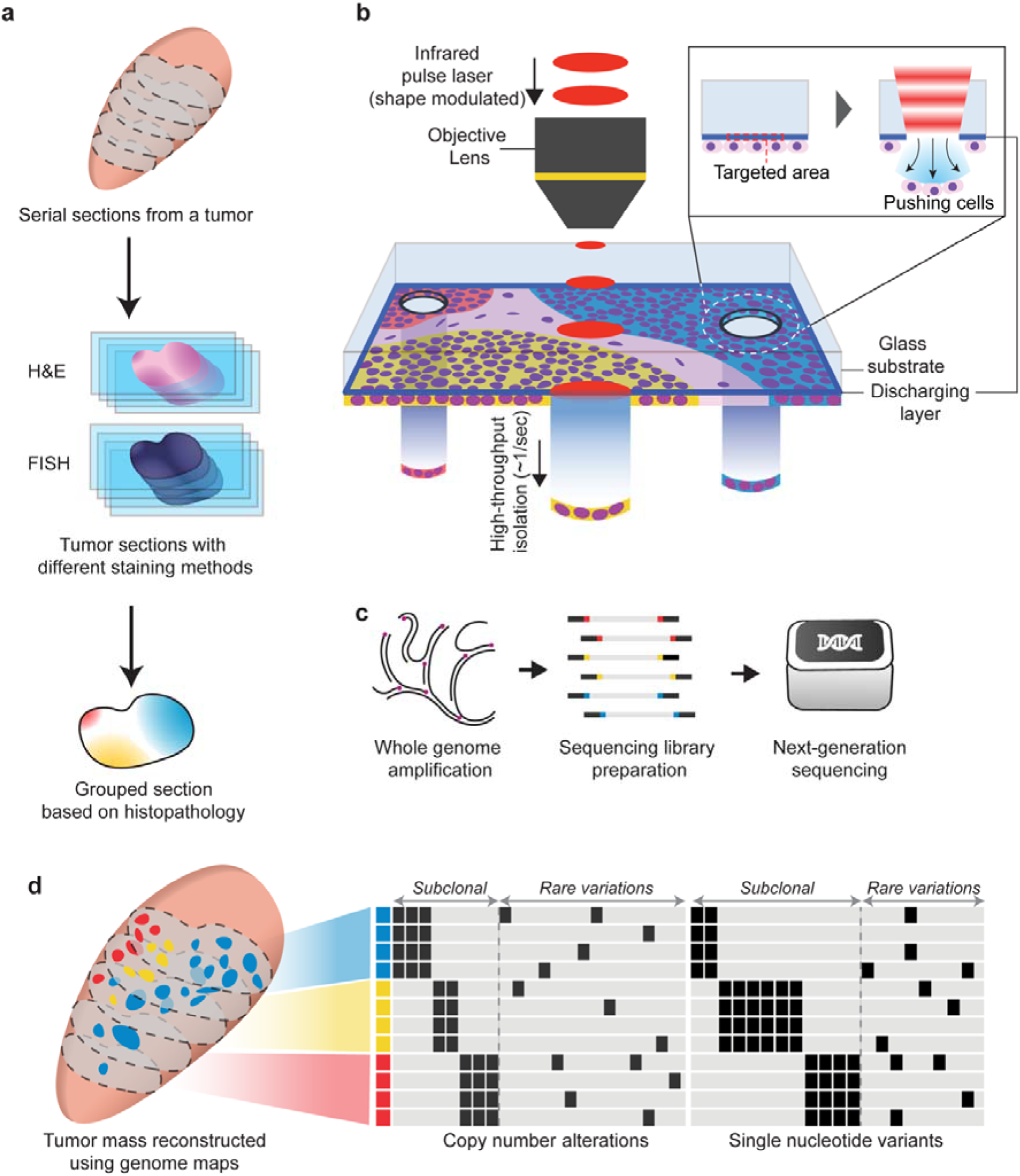
Phenotype-based high-throughput laser-aided isolation and sequencing (PHLI-seq) bridges the genotypic information to the corresponding phenotypic one in high-throughput. (**a**) The tumour mass is sectioned and stained with H&E. The section is prepared on an discharging layer coated glass. The tissue section is imaged, and the cancer cells in the section are grouped. Using grouping by phenotypes, cells that represent minor populations can be chosen. Grouping can be aided by pathologist manual observations. (**b**) A tissue section or cells prepared on the discharging layer can be isolated with an infra-red (1064 nm) laser. Because the discharging layer readily vaporizes where the infrared laser is applied, the cells on the vaporized area can be released downward by pressure due to vaporization. (**c** and **d**) After isolating the cells, we performed whole-genome amplification and massively parallel sequencing to analyse the CNA and SNV with their spatial organization in the tissue context.

After determining which cells to isolate, a single shot of near-infrared laser (1064 nm) was applied to each targeted area of the tissue. Because the discharging layer readily vaporized the ITO, to which the infrared laser was applied, the prepared cells could be transferred downward by the pressure (**Fig. 1b**). In contrast to laser microdissection using ultraviolet (UV) cutting and catapulting^9,25^, our method, which uses an infrared laser and one-shot isolation step, causes no damage to the nucleic acids (**Supplementary Fig. 1**) and can be performed rapidly, making it suitable for recovering high-quality DNA from single cells or small numbers of cells in a high-throughput manner. Moreover, our discharging method is advantageous over standard laser microdissection using an infrared laser to heat a thermoplastic film, which then expands and captures the cells of interest. Although this method is free of UV-induced damage, the thermoplastic film-based method requires physical contact between the film and the sample, and tearing off the cells by removing the cell-captured cap. This physical contact method causes inherent problems, such as limited throughput and scalability, debris or non-selected cell adhesion, and limited applicability depending on the sample condition^10,11^. Using PHLI-seq, we could isolate a region of cells or even single cells from various samples, regardless of how closely positioned were the single cells, while preserving information regarding the spatial and histopathological phenotype context in the tissue (Online Methods and **Supplementary Fig. 2**). Together with the operating software, PHLI-seq enables a high-throughput isolation of the targets (1 second per target) into the respective tubes (**Supplementary Note 1, Supplementary Fig. 3**, and **Supplementary Videos 1** and **2**). After the targets were retrieved, we used multiple displacement amplification (MDA) to amplify the whole genome. The amplified products were sequenced by massively parallel sequencing methods to analyse copy number alterations (CNA) and single nucleotide variants (SNV) with their spatial organization in the tissue context (**Fig. 1c** and **d**). To calculate the collection efficiency and the probability of obtaining high-quality sequencing data after cell isolation, we isolated singe-cell or very small numbers of cells from a cell line or an H&E-stained fresh frozen cancer tissue section. The collection efficiency was 92.5% (49/53) from single-cell isolations of the cell line sample (**Supplementary Fig. 4**). For the experiments with H&E-stained tissue sections, we thought that nucleic acids could be damaged for amplification by MDA during preservation, tissue sectioning, and staining. Therefore, we evaluated the probability of obtaining high-quality sequencing data from 5-cell and 1-cell isolations from H&E-stained tissue sections. We found that 81.3% (13/16) and 18.8% (3/16) of the 5-cell and 1-cell isolation experiments produced high genomic coverage and correct copy number profiles. Based on the sequencing data, we also found that high genomic coverage and a high correlation with the true copy number value of the whole-genome amplified samples were related to low C_T_ values for real-time monitoring of the amplification reactions (**Supplementary Fig. 4**). Therefore, we monitored the MDA procedure using a real-time PCR machine and validated the products by PCR using 16 primer panels to enrich the high-quality amplified products before sequencing (Online Methods and **Supplementary Table 1**). To confirm the sequencing performance, we isolated the HL60 cells using the PHLI-seq method, sequenced their genomes, and tested the coverage breadth, allele dropout (ADO), and false positive rate (FPR) (**Supplementary Note 2**, **Supplementary Fig. 5**, and **Supplementary Fig. 6**). Furthermore, we thoroughly tested whether the irradiating IR laser pulse and vaporizing discharging layer generate DNA fragmentations. However, we could not detect any signs of fragmentation in the sequencing data, even from cells that had repeatedly been irradiated by the IR laser pulse (**Supplementary Fig. 1**).

### Applying PHLI-seq to hormone receptor-positive/Human epidermal growth factor receptor 2-positive breast cancer and analysing copy number alterations

Next, we applied PHLI-seq to a hormone receptor (HR)-positive/ Human epidermal growth factor receptor 2(HER2)-positive invasive ductal carcinoma (IDC) to demonstrate that the genomes of the cell clusters from a stained tissue section can be effectively analysed. A tissue section of a preserved fresh frozen sample was prepared and stained with haematoxylin & eosin (H&E). The prepared section was scanned using an automated microscope to generate a high-resolution whole-section image (**Fig. 2a** and Online Methods). The image was segmented into cell clusters to generate a binary image, and spatial information and morphological phenotypes were extracted from each cell cluster. Then, a grouping of the cell clusters was performed using weighted hierarchical clustering to generate six groups (**Fig. 2b**, Online Methods, and **Supplementary Fig. 7**). Based on an additional inspection by pathologists, 53 cell clusters were selected from the groups for analysis by PHLI-seq (**Fig. 2c** and **d**). With an average of 20-30 cells in each cell cluster, whole-genome amplification and quality filtering were performed (Online Methods). The filter-passed samples were analysed by low-depth whole genome (n=53), whole exome (n=12), and targeted sequencing (n=53) (Online Methods).

**Figure 2.**
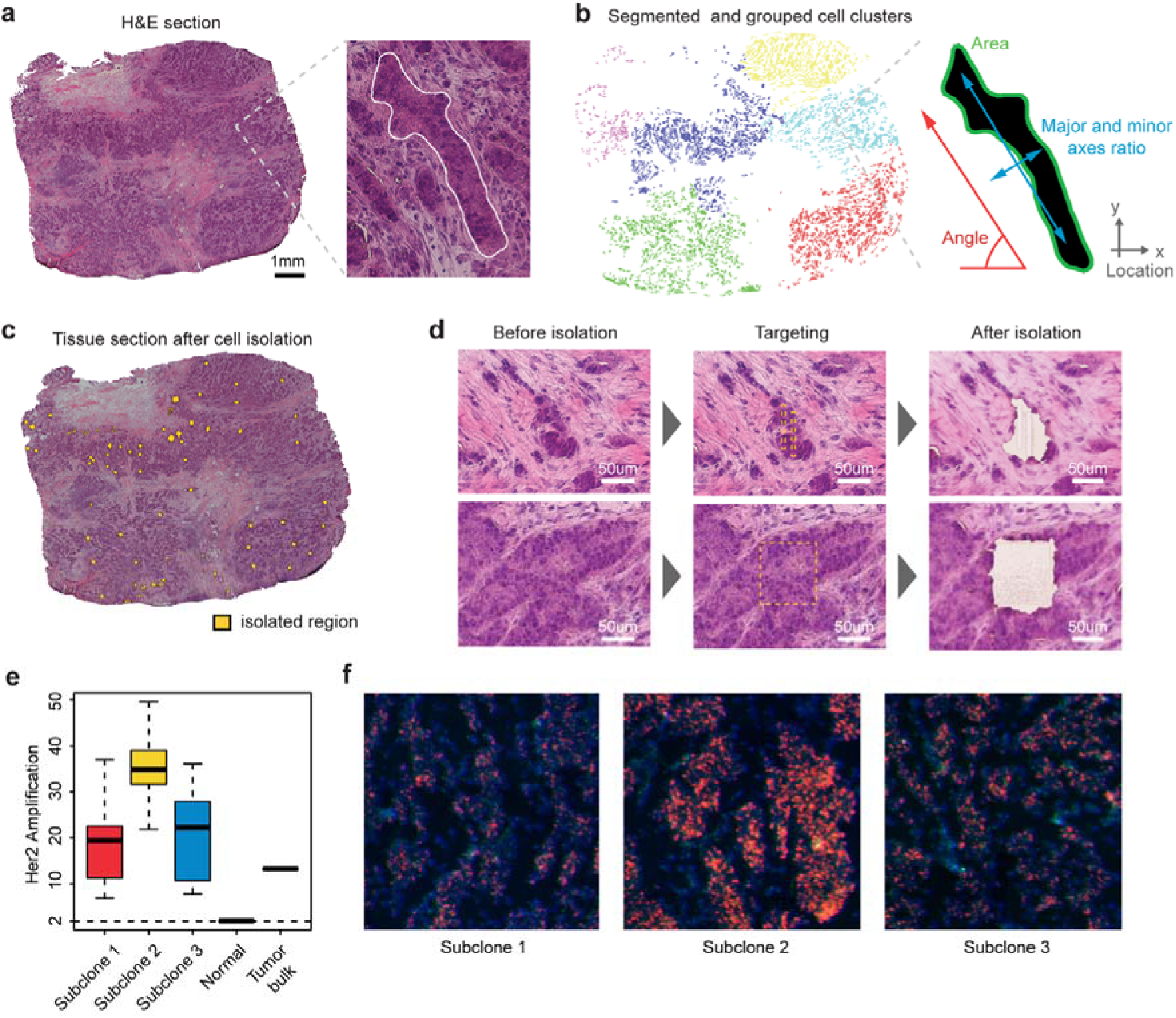
Fifty-three cell clusters were isolated and sequenced by PHLI-seq after grouping the cell clusters in HER2-positive breast cancer based on their location and morphology. (**a**) The H&E tissue section was imaged using a whole-slide imaging system. The enlarged image shows a typical example of a cell cluster. We highlighted the boundary of the cell cluster with a white solid line. (**b**) The H&E section image was segmented into cell clusters to generate the binary image. The cell cluster locations and three morphological features were extracted for grouping. Based on the information, the cell clusters were grouped by a weighted hierarchical clustering method to generate six groups, which are represented by six distinct colours. (**c**) Tissue section imaged after cell isolation. (**d**) Tissue images before and after isolation by the PHLI-seq system. The shape of the infra-red laser pulse can be modulated to isolate various shapes of cell clusters. (**e** and **f**) HER2 gene copy numbers in subclones identified by whole genome sequencing analysis and their respective FISH images.

First, genome-wide CNA analysis by low-depth whole-genome sequencing revealed three major subclonal populations in the tumour sample (**Fig. 3a** and **b**, approximate unbiased p-value > 0.99, multiscale bootstrap resampling with 10,000 iterations, Online Methods). The three subclonal populations had both shared and unique alteration profiles. The shared alterations include 1q gain, 8q gain, 8p loss and HER2 amplifications, all of which had been previously reported as frequent CNAs in human breast cancer and other types of cancer^26,27^. In this sample, the causative fitness gains underlying breast carcinogenesis appeared to take place in the *MYC* (8q24), *AKT3* (1q44), or *ERBB2* (HER2) (17q12) regions, all of which were gained or amplified in 12.4%∽21.3% of cases in The Cancer Genome Atlas (TCGA) project. Additionally, the up-regulation of these genes has been known to be associated with the causation of breast cancer^26^. In contrast, subclones 1, 2 and 3 harbour unique gains or focal amplifications at 6q22.32-q23.2, 2q14.1-q14.2 and 8p21.3-p22, respectively. Among them, the amplification of *CUG2* (Cancer-Upregulated Gene 2, also called *CENPW*, 6q22.32), which is known to be associated with tumour progression^28^, might contribute to the high proliferative ability observed in subclone 1. One interesting observation is that the CNAs status was clearly divided into three distinct populations with no intermediate subclones. Since intermediate subclones might be excluded from the sampling process, we isolated additional cell clusters (n=27) at the boundaries between subclones. The isolated samples were analysed by low-depth whole-genome sequencing. Then, clustering analysis was performed based on the inferred copy number data for both previously isolated (n=53) and additionally isolated samples (n=27). The results showed that the 80 cell clusters from the HER2-positive tissue sections were classified into one of the three previously defined cancer subclones (**Supplementary Fig. 8**). This result reinforces evidence for the punctuated copy number change followed by a period of stasis, as demonstrated in previous studies^16,18,29^.

**Figure 3.**
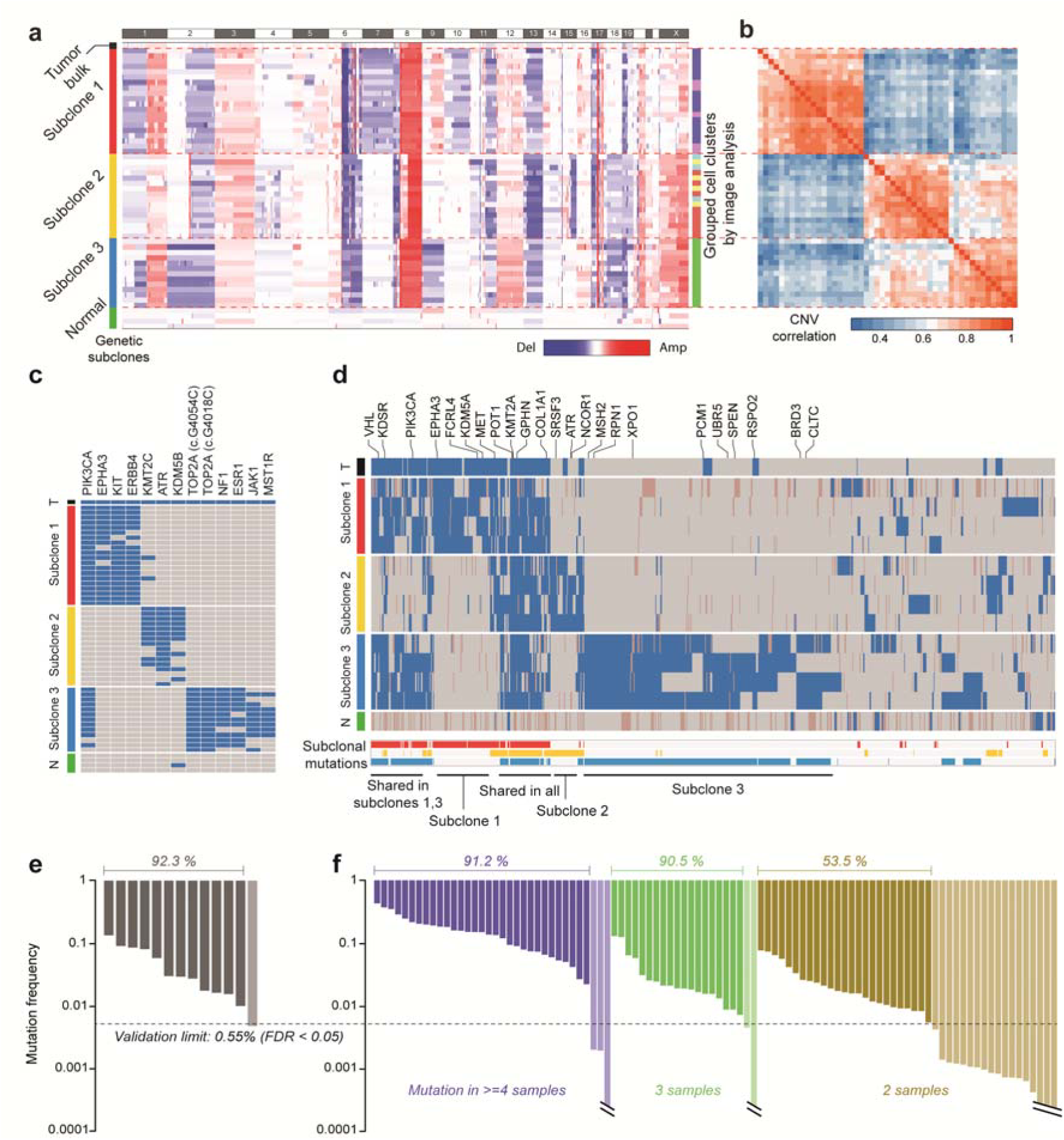
CNA and SNV analysis by whole-genome, targeted, and whole-exome sequencing and its validation by single-molecule deep sequencing reveal three different subclones of the cancer tissue. (**a**) After performing low-depth whole-genome sequencing (0.16 Gb/sample), CNA were determined as described in the Online Methods. Rows were reordered to cluster samples by the correlation of their CNAs. Excluding the tumour and normal bulk, we found three distinct genetic subclones. The three subclones had both shared and exclusive CNA events. (**b**) The correlation matrix of the copy number data also produced the three subclones. Unsupervised clustering was used for reordering of the samples. (**c**) Targeted sequencing of the 53 isolated cell clusters. As in the CNA analysis, we found three distinct genetic subclones. The three subclones had unique somatic mutation patterns. (**d**) Whole-exome sequencing of 4 samples from each subclone. Subclones 1 and 3 shared a large portion of mutations, including the *PIK3CA* mutation. (**e** and **f**) Mutation validation using single-molecule deep sequencing. We set our validation limit to 0.55%, which limited the false discovery rate to < 0.05.

### Whole-exome and Targeted sequencing to analyse sequence alterations

To investigate somatic SNV, we performed targeted sequencing of 121 genes associated with breast cancer (Online Methods and **Supplementary Table 2**). The results revealed unique mutational profiles in each subclone, consistent with those determined by whole-genome sequencing (**Fig. 3c**). In our targeted sequencing analysis of 53 cell cluster samples (20 (subclone 1), 16 (subclone 2), 13 (subclone 3), and 4 (no classification)), we found that mutations in *PIK3CA, EPHA3, KIT, ERBB4* and *KMT2C* occurred in subclone 1, mutations in *KMT2C, ATR* and *KDM5B* in subclone 2, and mutations in *TOP2A, NF1, ESR1, JAK1* and *MST1R* in subclone 3. Notably, PIK3CA p.M1043I, which has previously been known to be an oncogenic mutation that causes an increase in PI3K lipid kinase activity, constitutive AKT activation and the transformation of NIH3T3 cells and chick embryo fibroblasts, was shared between subclones 1 and 3^30,31^. In addition, PIK3CA p.M1043I and EPHA3 p.E794K in subclone 1 and ATR p.R2363X in subclone 2 have been reported to be recurrently observed in breast and other types of cancer^27,32^. Intriguingly, the stop-gain mutation ATR p.R2363X destroyed the PI3_PI4_kinase domain, a key functional structure, in the tumour suppressor ATR, which plays a role in the DNA mismatch repair pathway. Another validated mutation that could cooperate in driving the subclonal heterogeneity is ESR1 p.E380Q in subclone 3, which has been known to be associated with endocrine therapy resistance in a breast cancer xenograft model^33^. Although the nonsynonymous EPHA3 p.E794K is a novel mutation that has not been reported to date in dbSNP, we confirmed that this mutation with the highest deleterious risk scores (SIFT_score, 0 and Polyphen2_HDIV_score, 1) occurred in the protein tyrosine kinase domain (PF07714) of EPHA3, meriting further functional analysis in the future. The stop-gain mutation KMT2C p.Q3417X in subclone 1 completely truncated the SET domain (PF00856), a key functional structure, implying a consistency with a recent report showing that low KMT2C expression is associated with a poor outcome in ER-positive breast cancer patients^34^. In cooperation with the oncogenic copy number amplifications mentioned above, those deleterious mutations in the three subclones may contribute significantly to the independent oncogenic evolution of each subclone.

For further analysis, we performed whole-exome sequencing of four samples selected from each subclone (**Fig. 3d**). We found that 75 mutations were shared in the three subclones and that 99, 75, and 382 mutations in *VHL, KDSR, PIK3CA, EPHA3, FCRL4, KDM5A, MET, POT1, KMT2A, GPHN, COL1A1, SRSF3, ATR, NCOR1, MSH2, RPN1*, and *XPO1* occurred exclusively in subclones 1, 2, and 3, respectively. In contrast to the whole-exome mutation profiles in the three subclones by PHLI-seq, we could not find such representative mutation profiles in the sequencing data from the tumour bulk. The mutations detected in the tumour bulk mainly covered the mutational profile in subclone 1 (78.8% of exclusive mutations), whereas only approximately 6.81% of the mutations in subclone 3 were detected in the tumour bulk (**Supplementary Fig. 9**). This result implies that PHLI-seq can provide rich information about subclonality and variants with a low-level allele fraction in heterogeneous tumours, even those with subclones that are too minor to be detected by conventional methods.

Next, we validated our SNV analysis by single-molecule deep sequencing. We randomly selected a portion of mutations that were detected using PHLI-seq and generated a sequencing library for the targeted sites by tagging a unique molecular barcode to each DNA molecule to precisely discriminate the next-generation sequencing results from errors (Online Methods; **Supplementary Fig. 10**). Sequencing was performed to read the targeted region with an 18,182-fold single-molecule read depth. We set our validation limit to 0.55% to limit the false discovery rate to < 0.05. The results validated 92.3% (12/13) of the subclonal mutations detected by targeted sequencing (**Fig. 3e**). Moreover, 72.2% (78/108) of the mutations observed in the whole-exome sequencing were validated. The validation rate was 90.9% (50/55) when mutations that were observed in more than three samples were considered (**Fig. 3f**).

### Evolution of tumour subclones and their spatial organization in the tissue context

Based on the CNA and SNV analyses, we could infer the evolutionary history of the subclones in the tumour (**Fig. 4a**). An ancestral precursor clone could be generated by gaining an advantage for tumorigenic proliferation by accumulating the CNAs and mutations shared by the major three subclones. However, we cannot rule out the possibility that the genesis of CNAs and mutations shared by the three subclones might have been completed over several generations of previous initial subclones originating from a *bona fide* ancestor clone, giving rise to the birth of a plausible precursor clone that harboured the variants shared by the three subclones. Then, one cell in the initial precursor clone could have acquired other variants, becoming a subsequent precursor clone for the generation of subclones 1 and 3. Other cells in the initial precursor clone could accumulate different variants that could consequently give rise to subclone 2, evolutionarily branching out from subclones 1 and 3. This result can be inferred from the observation that subclones 1 and 3 have many common mutations, including PIK3CA p.M1043I, whereas subclone 2 has exclusive mutations and mutations that are common to all three subclones (**Fig. 4a**). Moreover, a copy number analysis of the *HER2* gene showed that subclone 2 has a much higher *HER2* amplification level than subclones 1 and 3, further supporting this scenario (**Fig. 2e** and **f**). After being split into the two lineages, the precursor subclones would have accumulated copy number changes or mutations independently, and one of the two could be further split to ultimately give rise to subclones 1 and 3. Subclone 3 could have accumulated more mutations, potentially because of the exclusive mutation in the MSH2 gene encoding a component of the post-replicative DNA mismatch repair system.

**Figure 4.**
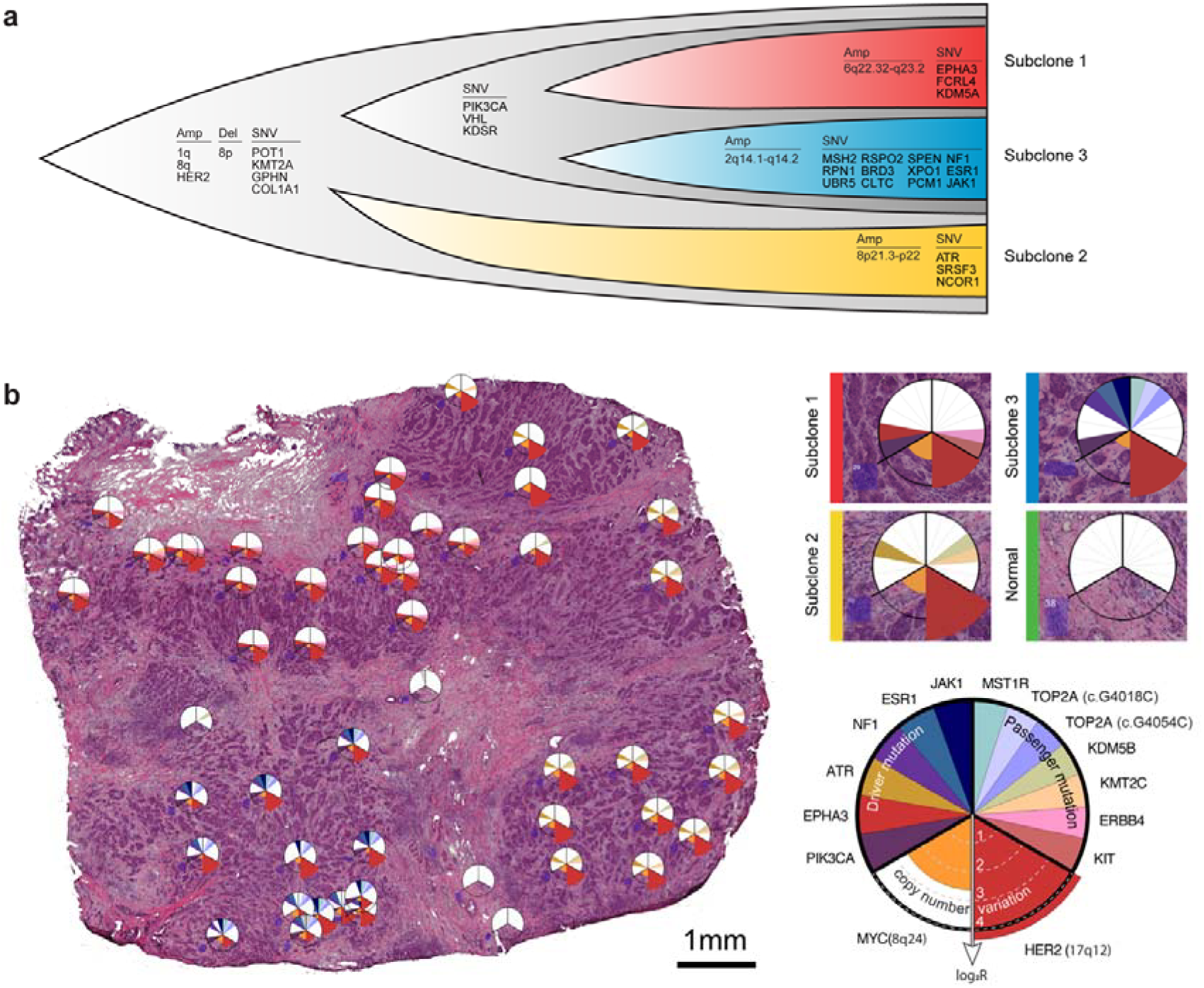
The evolution of the three subclones and their spatial organization is inferred through PHLI-seq. (**a**) Alterations shared by the three subclones may have led to the early stage of tumorigenesis. Then, subclone 2 and the ancestor of subclones 1 and 3 may have divided and accumulated unshared alterations. (**b**) Spatial mapping of genomic data showing that each subclone is spatially segregated, with stroma between each subclone.

Finally, we mapped the detailed information for the CNAs, driver mutations, and passenger mutations to the topological information and spatial positions of the tumour tissue at 10 µm resolution (**Fig. 4b**). The three subclones were found to be spatially segregated in the tumour mass. As shown in Fig. 4b, whereas the heterogeneity of the tumour tissue is clear from the detection of the three different subclones, the micro area occupied by each subclone exhibits no mingling with cells from other subclones. This finding implies that the three subclones are independent with well-established tumorigenic advantages and strongly suggests that a combination of diverse drugs for inhibiting different subclones in each patient should be a future therapeutic strategy for personalized cancer medicine.

### Constructing and visualizing a cancer genomic map in a three-dimensional spatial context

We further analysed consecutive sections of a triple-negative (estrogen/progesterone receptor and HER2-negative) breast cancer sample to discover how heterogeneous tumour subclones exist in the three-dimensional space of the tissue and to demonstrate how PHLI-seq can be an empowering tool to bridge genomics to histopathology (**Fig. 5a**). The size of the tumour was about 7 × 6 × 5mm, and seven tissue slices with an interval of 700 µm between each of them were used to prepare H&E sections for PHLI-seq. A total of 177 cell clusters from the seven H&E sections were isolated and sequenced by PHLI-seq as described. Before the isolation, cancer cells with various phenotypes were identified by histopathological evaluation from H&E and IHC (AR, CK5/6, Ki-67, and p53; **Supplementary Fig. 13**) sections. Based on the phenotypic information, the 177 cell clusters were selected to discover genetic heterogeneity in tumour cells with various phenotypes. After isolating the targeted cell clusters, we performed low-depth whole-genome sequencing to obtain an image of the heterogeneity of CNAs (**Fig. 5b**). We discovered three genetic subclones, and they were denoted *in situ* clone 1, *in situ* clone 2, and invasive clone. *In situ* clone 1 and *in situ* clone 2 had the same copy number profiles, except for the deletion of a q arm of chromosome 16 and a p arm of chromosome 17 in *in situ* clone 2, suggesting that the *in situ* clone 2 may have derived from ancestral cells of the *in situ* clone 1 by additional chromosomal deletions. The invasive clone showed additional chromosomal amplification in chromosome 1 (q arm), 6, 7, 8, 10, and X. To investigate more genomic differences at the single nucleotide level, we performed whole exome sequencing for 11 clusters in the three clones (**Fig 5c**). The result shows that the tumour cells in the invasive clone had mutations which do not exist in *in situ* clones. Surprisingly, even *in situ* clones had mutations which were not observed in the invasive clone. This may suggest that IDC is derived from an early ancestry of ductal carcinoma *in situ* (DCIS), atypical ductal hyperplasia (ADH), or other benign cells, not directly from DCIS in a linear manner. Moreover, *in situ* clone 2 showed mutations exclusive to *in situ* clone 1, which supports the previous explanation that *in situ* clone 2 may be derived from the ancestral cells of *in situ* clone 1, based on CNA profiles.

**Figure 5.**
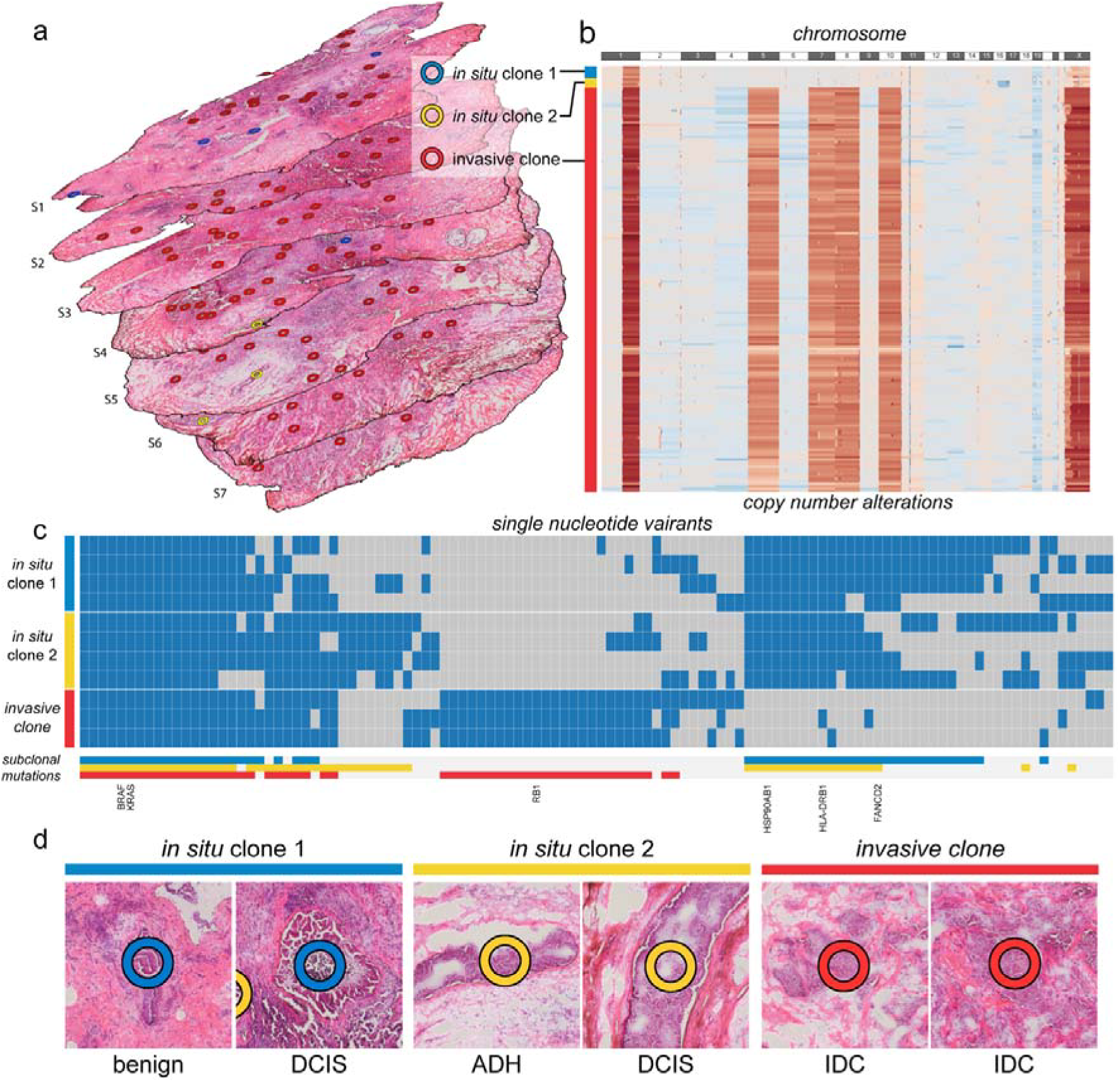
The 3-dimensional tumour mass was investigated using PHLI-seq. (**a**) A total of 177 cell clusters were isolated and analysed from 7 consecutive tissue sections from a triple-negative breast tumour. Before the isolation, cancer cells with various phenotypes were identified by histopathological evaluation from H&E and IHC (AR, CK5/6, Ki-67, and p53; **Supplementary Fig. 13**) sections. Based on the phenotypic information, the 177 cell clusters were selected to discover genetic heterogeneity in tumour cells with various phenotypes. Seven consecutive tissue sections had a 700-µm interval between each section. (**b**) Whole-genome sequencing of the 177 cell clusters in the tumour discovered three subclones. The *in situ* clones 1 and 2 shared CNAs in chromosomes 1 and X. On the other hand, the invasive clone had considerably more amplifications in chromosome 1 and X, and additional amplifications in chromosome 5, 7, 8, and 10. (**c**) Whole-exome sequencing of the selected samples from each subclone. The subclonal mutations are labelled in the corresponding colour to demonstrate shared and unique mutations. (**d**) *In situ* clone 1 included DCIS and benign usual ductal hyperplasia, whereas *in situ* clone 2 included DCIS and ADH. Histopathologic evaluation of the invasive clone showed that every cell clusters in this subclone were IDC.

Within the two *in situ* clones determined by whole genome sequencing, cells with different histopathologic features were observed. DCIS and benign usual ductal hyperplasia included in *in situ* clone 1, whereas DCIS and ADH were included in *in situ* clone 2. Histopathologic evaluation of the invasive clone showed that every cell clusters in the invasive clone were IDC (**Fig. 5d**). A three-dimensional reconstruction of the sections allowed identification of *in situ* clone 2 clusters observed in a vicinity in consecutive sections S4, S5, S7, and S7, suggesting that combined analysis of histopathology and spatially resolved genomics enabled by PHLI-seq has potential to contribute to clinical diagnostics.

## DISCUSSION

In this study, we developed a new technology that effectively discovers the genetic heterogeneity of tumours with accurate mapping of genomic alterations to the spatial information of the tissues. We identified three subclones in a HR-positive/HER2-positive breast cancer tissue and discovered subclonal CNAs and point mutations. Here, we found that the majority of CNAs are shared by all groups, whereas those of SNVs are not. Similar to a prior report^12^, this result indicates that most chromosomal rearrangements preceded the generation of point mutations. Moreover, when only CNAs are considered, the evolution of the tumour seems to have followed the model of punctuated evolution with a one-time event, rather than the model of gradual evolution^16,18^. However, regarding the SNVs, the evolution cannot be explained by the model of punctuated evolution with a one-time event. Because subclones 1 and 3 share many SNVs, the divergence of subclones 1 and 3 from the ancestral population seems to have occurred later than the evolutionary burst that generated subclone 2 and the ancestral population of subclones 1 and 3. Therefore, considering both the CNAs and the SNVs, our data can be explained by the model of punctuated copy number evolution with two-time events. Overall, we visualized the three genetic subclones of the HR-positive/HER2-positive breast cancer in tissue context with their genetic history of punctuated copy number evolution.

We also applied PHLI-seq to serial breast cancer tissue sections to construct and visualize a cancer genomic map in a three-dimensional spatial context (**Supplementary Video 3**). From 177 cell clusters in a 7 mm × 6 mm × 5 mm space, we identified three genetic subclones encompassing the histopathologically classified benign cancer cells ADH, DCIS, and IDC. We performed CNA and SNV analyses by WES to elucidate the evolutionary history of the cancer cells in relation to the histopathological information. In three-dimensional space, the invasive clone was located throughout the entire tissue, while the *in situ* clones were confined to relatively smaller areas. Although *in situ* clone 2 clusters were observed in a vicinity in four consecutive sections, it is interesting that the *in situ* clone 1 was observed in sections S1 and S4, but not in between them (**Fig. 5a**). Despite thorough observation of sections S2 and S3, we could not detect any benign cells or DCIS in the histopathological evaluation. Moreover, we could not detect exact spatial linkage between *in situ* clones 1 (sections S1 and S4) and 2 (sections S4, S5, S6, and S7). One explanation could be that the invasiveness of tumour cells in the invasive clone resulted in the spatial separation of the *in situ* clones, especially between the *in situ* clone 1 cell clusters in sections S1 and S4. This suggests that different tumours show various and complex spatial context, and PHLI-seq is the only method that allows for correlated analysis of sequencing data and histopathological spatial features.

This new high-throughput technology can accurately bridge the histopathological and genomic alteration landscape by leveraging the map of genomic alteration at the single-cell level to spatial positions in a tissue, allowing an unprecedented better understanding of carcinogenesis. Additionally, by comparing primary cancers with metastatic or recurrent cancers by high-precision analysis of micro-local tissue areas at the single-cell level using PHLI-seq, one can discover novel important molecular features concerning the carcinogenic transition between different tumorigenic stages. In addition, this technology can provide novel insights at the high-precision single-cell level of interactions between a tumour cell and its microenvironment, which has not been clearly elucidated by previous conventional techniques.

Another new innovative improvement supplied by this technology is its exploitation of different staining modalities other than H&E, FISH, and IHC, among others. Additionally, PHLI-seq can incorporate machine learning to integrate histopathological information, three-dimensional positions, and genomic alteration landscapes, which will elucidate the relationship between histopathology and genomic and molecular features that have remained obscure based on conventional methods alone. PHLI-seq will make a significant contribution to future subclonal evolutionary analyses of carcinogenesis and the discovery of novel therapeutic targets at the oncogenic subclonal level.

Previously, several groups have developed computational analytic methods for H&E-stained images based on the single nucleus morphology^35,36^, although we utilized image analysis techniques for a different purpose than those of Yuan et al. and Beck et al. We developed a method for image-based grouping of “potential” genetic subclones of cancer cells by grouping cell clusters in an H&E section based on their location and morphology. With the guide of the grouping, we performed PHLI-seq to discover the genetic heterogeneity of the cancer cells. For this purpose, cell cluster-based image analysis would be more suitable than approaches based on single-cell nucleus morphology (Online Methods). However, a limitation of our method is that we did not select the four features based on a large dataset. Yuan et al. and Beck et al. used hundreds of H&E-stained images to establish classifiers based on hundreds of features. As our goal was to develop a technique to reveal subclonal heterogeneity among cancer cells that may have similar cellular phenotype traits, we could not build the image analysis pipeline based on many samples. However, if such a technique can be applied, PHLI-seq would be much more effective. We believe that cutting-edge histopathological image analysis and 3D image visualization techniques can greatly improve the potential and utility of PHLI-seq in the future.

From a clinical perspective, PHLI-seq can be applied to broader areas. As mentioned earlier, genomic regions showing copy number amplifications shared by the three subclones harbour the loci of the oncogenes *MYC, ERBB2* (HER2) and *AKT*. In addition, another oncogenic driver mutations are present in subclone 1 (*PIK3CA* and *EPHA3*), subclone 2 (*ATR*), and subclone 3 (*NF1* and *ESR1*)^46,47,48^. For example, for the subclonal tumorigenic heterogeneity status identified by PHLI-seq in this study, the following insight for a combined therapeutic drug treatment strategy may be proposed. First, to inhibit a background tumour from which the three subclones originated, HER2 inhibitor, MYC inhibitor and AKT inhibitor might be used. Second, to specifically inhibit subclone 1, PI3K/AKT/mTOR inhibitors and agonistic anti-EPHA3 mAb IIIA4 might be used, given the suppression of the background tumorigenic activity. Third, to prevent subclone 3 specifically, AZD9496 (an oral oestrogen receptor inhibitor) might be used. Finally, small molecule ATR inhibitors (ATRi) might be used to specifically prevent subclone 2. As some of the above-mentioned drugs are now in nearly final stages of their clinical trials, they may be in use in upcoming years pending FDA approvals. Although such a combined therapeutic strategy could not be applied to patients because our major concern in this study was to develop the novel PHLI-seq approach and the proper timing for its application in patients was lost, we propose that PHLI-seq can be applied to a variety of cancer types in the future to provide insights on subclonal tumorigenic heterogeneity for establishing combined pharmaceutic and treatment strategies, similar to the above-mentioned example.

Finally, it should be noted that spatial genetic information can affect clinical interpretations because cancer cells evolve through various geographic conditions and microenvironments in the tissue and can act differently depending on their tumour location. Thus, the PHLI-seq platform meets the needs of cutting-edge cancer biology, which links histopathology to genomics to enable a synergistic and more precise interpretation of cancer.

## ONLINE METHODS

### Sample Preparation

Human fresh-frozen breast cancer tissues were obtained from the Department of Surgery, Seoul National University Hospital, and analysed under the approved Institutional Review Board (IRB) protocol (IRB No. 1207-119-420). Frozen breast cancer tissues were stored at −80 °C until they were sliced into 10-µm-thick sections using a Leica CM3050 cryostat (Leica Microsystems GmbH, Wetzlar, Germany). Tissue sections were thaw-mounted onto ITO-coated glass slides. Glass slides were stored at −20 °C until analysis. The tissue sections were dried for 15 minutes at room temperature and subjected to a modified haematoxylin and eosin (H&E) staining technique according to the following protocol: 1) rinse in tap water for 5 minutes, 2) stain in Harris haematoxylin solution (Merck, Darmstadt, Germany) for 3 minutes, 3) rinse in tap water (quick dip), 4) rinse in 1% HCl solution/EtOH (quick dip), 5) rinse in tap water for 5 minutes, 6) counterstain with eosin Y (BBC Biochemical, Mount Vernon, WA) for 3 seconds, 7) rinse in water (10 dips), 8) dehydrate in 70% EtOH (10 dips), 9) dehydrate in 90% (10 dips), and 10) in 100% EtOH (10 dips). Sections were allowed to dry at room temperature.

The tissue used in the 2D experiment was obtained from a 57-year-old woman who underwent total mastectomy with axillary lymph node dissection in April 2014. The breast cancer was a Stage IIIC (pT2N3M0, AJCC 7^th^ TNM Staging) IDC which was positive for estrogen receptor (95%), progesterone receptor (2%), and HER-2 (+++/3) on IHC, as evaluated according to the American Society of Clinical Oncology and College of American Pathologists (ASCO/CAP) guidelines. The tissue used for 3D mapping was obtained from a 56-year-old woman who underwent Total mastectomy with axillary lymph node dissection in September 2015. The breast cancer was a Stage IIA (pT2N0M0, AJCC 7^th^ TNM Staging) IDC which was negative for estrogen receptor (1%), progesterone receptor (negative) and HER-2 (-/3) on IHC.

### Whole-slide imaging to produce digital slides

For whole-slide imaging, a slide was removed from the refrigerator and left at room temperature for 10 minutes. After placing the slide on an automated microscope (Inverted Microscope Eclipse Ti-E, Nikon Instruments Inc., Melville, NY), we set the scanning region of the slide. Scanning was performed using a 20x lens, and stitching was carried out by an internal algorithm (NIS-Elements AR Auto Research, Nikon Instruments Inc., Melville, NY).

### Grouping cell clusters based on spatial and phenotypic information

To group the cell clusters based on spatial and phenotypic information, we performed the following steps: 1) whole-slide imaging, 2) cell cluster segmentation, 3) extraction of spatial and phenotypic information, and 4) weighted hierarchical clustering. 1) We imaged the whole slide as described in the section ‘Whole-slide imaging to produce digital slides’. 2) Cell cluster segmentation was carried out using a conditional random field. The whole-slide image was split into small images of 1024 × 1024 pixels. The learning set was constructed for 50 split images. Then, learning and segmentation were performed using the conditional random field^37^. After segmentation, split images were merged into one image of the original size. 3) We indexed each cell cluster in the merged image using the connected components. Then, four features were extracted for each cell cluster: ‘position’, ‘cluster area’, ‘major axis over minor axis’, and ‘angle’. ‘Position’ is the centroid of the cell cluster. ‘Cluster area’ is the number of pixels of the cell cluster. ‘Major over minor axis’ is a ratio of the major axis over the minor axis, which estimates how the ellipse-like figure fitted to the cell cluster is. ‘Angle’ is the angle between the horizontal line and a straight line parallel to longer side of the rectangle, which estimates the tilt bias of the ellipse-like figure from the vertical line. All scripts were written in C++ and using the OpenCV library for connected components and other features. 4) We performed weighted hierarchical clustering using the four features to find the optimal clustered groups. To find the optimal groups, we maximized a score as follows:

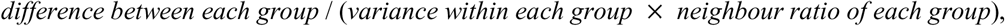

where

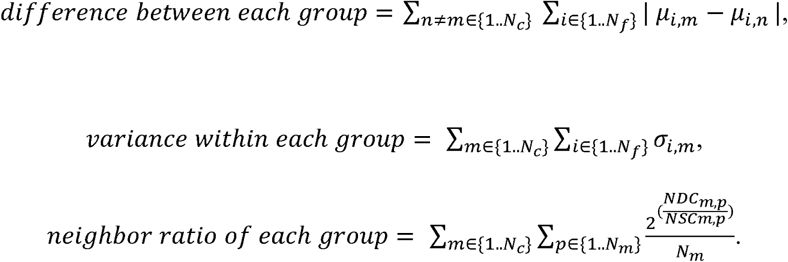

Here, *µ*_*i,m*_ and *a*_*i,m*_ represent the mean and the standard deviation of the *i*^*th*^ feature in the *m*^*th*^ group, *N*_*c*_ and *N*_*f*_ are the number of groups and features, p is the index of a cell cluster and *N*_*m*_ is the number of cell clusters in the *m*^*th*^ group. *NSC*_*m,p*_ is the number of cell clusters that are classified into the *m*^*th*^ group among the ten nearest neighboring cell clusters to the *p*^*th*^ cell cluster of the *m*^*th*^ group. In contrast, *NDC*_*m,p*_ is the number of cell clusters that are not classified into the *m*^*th*^ group among the ten nearest neighboring cell clusters to the *p*^*th*^ cell cluster of the *m*^*th*^ group. We maximized the score by fixing the weight for ‘position’ as one, sweeping the weight of the other features from 0.1 to 10 with an increment of 0.25. All scripts were written in R using the stats library for hierarchical clustering.

Previously, several groups have developed computational analytic methods of H&E images based on single nuclear morphology^35,36^. Unlike these approaches, which utilize a single-cell nucleus as a base unit for classifying cell types, we have used the ‘cell cluster’ as the basis for calculations for several reasons. First, we think that the cell types (e.g., cancer, stromal, and immune cells) can be determined based on the single nucleus morphology because their biological functions and phenotypes clearly differ from each other. However, it is hard to predict ‘genetic’ subclones in cancer cells using the single nucleus image because they are all cancer cells and have relatively similar properties (e.g., large and rounded nuclei) in many cases. Therefore, we focused on parameters that can reflect the collective functions of each subclone, such as the growth level, growth direction, and stromal infiltration, which can differ for each subclone. These phenomena are reflected in the morphology of the cell cluster rather than the single nuclei. Second, because a single-cell in a tissue section may not contain an intact nucleus, we needed to isolate a cell cluster or a small number of cells in a cell cluster to read the full genomic information. Although we sectioned tissues to a thickness of 10 µm, some cells in the section may have a partial nucleus. We considered it to be a poorly controlled approach to isolate and analyse a single-cell in a tissue section. Therefore, we decided to use a cell cluster containing a small number of cells for the sequence analysis. Consequently, we used a cell cluster for image analysis because it was our base unit of sequence analysis. Based on discussions with a pathologist, we selected four features: position, cluster area, major axis over minor axis, and angle. The position and cluster area reflect the relative location of a subclone in a tissue and its growth level, respectively. The major axis over the minor axis is associated with invasiveness and stromal infiltration. Finally, the angle represents the growing direction of a subclone.

### PHLI-seq instrument

The PHLI-seq instrument comprises two motorized stages, a CCD camera, light source, laser source, pulse slit, objective lenses, and fluorescence modules **(Supplementary Fig. 3).** The two motorized stages (ACS Motion Control, Migdal, Israel) can be controlled automatically by communicating with a computer. One is for loading sample slides, and the other is for loading tubes to receive isolated cells. The CCD camera (Jenoptik, Jena, Germany) is installed to observe where the laser pulse will be applied through the objective lenses. A Nd:YAG nanosecond laser was purchased from Continuum (Minilite^TM^ Series ML II; Continuum, San Jose, CA). A slit is located in the light path between the laser source and the objective lens to control the region to be isolated. The slit is controlled either manually or automatically to adjust the size of the laser pulse. Objective lenses with various magnifications were purchased from Mitutoyo. The long working distance allows more space between the lens and the sample for user convenience.

### Software to automate the PHLI-seq procedure

We designed two different pieces of software, which were written in Python scripts (**Supplementary Scripts**) and available at Github (https://github.com/BiNEL-SNU/PHLI-seq). The first was built for the user (a pathologist in our case) to select the cells to be isolated. We shared the whole slide image with the user through a server, and the user ran the software to select the cells of interest while navigating the tissue image through the graphical user interface. After selection, the program produces two files: a text file with locational information about the region of interest, and the image file with the selected targets overlaid with transparent blue on the original image. Both files are required for the automated isolation of the target cells. The second software enables the automatic control of the PHLI-seq instrument. With this software, the users are able to control the slits, change the objective lenses, and move the motorized stages. It also enables automatic target isolation when two files from the first software are loaded. All tissue samples were isolated using an automatic function, while the cell line experiments were performed manually.

### Cell isolation and whole-genome amplification

Cells were isolated from tissue sections, cell lines, or blood smears that were spread on the ITO glass (Fine Chemicals Industry, Seoul, Korea), where ITO was coated on a glass by sputter deposition. An infrared laser was applied to the target area, vaporizing the ITO layer and discharging the target cells in the region. We used glass slides with a 100-nm-thick ITO layer. We tried coatings of 100 nm, 150 nm and 300 nm ITO layers for the experiments. However, there was no detectable difference between them. Cancer cells could be isolated from 4-10-µm-thick tissue sections at all thicknesses. The thickness of the ITO layer can affect the maximum tissue thickness at which the cells can be detached, although we did not undertake a thorough assessment. To separate thicker or harder tissues than in the present study, it may be better to use a thicker ITO layer, although we had no problem separating the cells from the bone tissue when using a 100-nm-thick ITO layer. Using an infrared laser in conjunction with the ITO discharging layer, we could accomplish cell isolation without damaging the cell (**Supplementary Fig. 11** and **Supplementary Fig. 12**). The 8-strip PCR tube caps for the retrieval of cells were pre-exposed under O_2_ plasma for 30 seconds. The cells were lysed using proteinase K (cat no. P4850-1ML, Sigma Aldrich) according to the manufacturer’s directions after the PCR tubes were centrifuged. For whole-genome amplification by multiple displacement amplification, we used GE’s Illustra Genomiphi V2 DNA amplification kit (cat no. 25-6600-30). We added 0.2 µl of SYBR green I (Life Technologies) into the reaction solution for real-time monitoring of the amplification. All amplified products were purified using Beckman Coulter’s Agencourt AMPure XP kit (cat no. A63880) immediately following the amplification reaction. To validate that the samples were thoroughly amplified, we used real-time whole-genome amplification monitoring and PCR validation with in-house designed 16-region primer panels (**Supplementary Table 1**). To discard poorly amplified samples, we used an MDA product that exhibited an observable amplification level within 40 minutes after the start of the reaction. Most of the amplified products yielded more than 1 μg, and 800 ng was used for Illumina library construction. To prevent carry-over contamination, the pipette tip, PCR tube, and cap for the reaction were stored in a clean bench equipped with UV light and treated with O_2_ plasma for 30 seconds before use. Additionally, we monitored the real-time amplification of non-template controls to ensure that no contaminants were transferred.

### Sequencing of the amplified genome

The whole-genome amplified products or genomic DNA extracts were fragmented using an EpiSonic Multi-Functional Bioprocessor 1100 (Epigentek) to generate a 150∽250-bp fragment distribution. The fragmented products underwent Illumina library preparation for end repair, 3’dA-tailing, adaptor ligation, and PCR amplification according to the manufacturers’ instructions. We used the Celemics NGS Library Preparation Kit (LI1096, Celemics, Seoul, Korea) for the whole-genome sequencing library preparation, SureSelectXT (Agilent, CA, US) for whole-exome sequencing, and the Celemics Customized Target Enrichment Kit (SICT96, Celemics, Seoul, Korea) for targeted sequencing. DNA purification was performed by ^TOPQ^XSEP MagBead (XB6050, Celemics, Seoul, Korea), and DNA libraries were amplified using the KAPA Library Amplification Kit (KAPA Biosystems, KK2602). Finally, the products were quantified by TapeStation 2200 (Agilent, CA, US). We used a HiSeq 2500 50SE (Illumina) to generate 0.16 Gb/sample for whole-genome sequencing and a HiSeq 2500 150PE (Illumina) to generate 5 G/sample and 0.88G/sample for whole-exome and targeted sequencing, respectively.

### Targeted sequencing panel

We chose 121 genes for the SNUH BCC (Seoul National University Hospital Breast Care Center Panel) based on the following criteria.

In our previous study, we performed whole-exome sequencing and RNA-Seq of 200 pairs of matched clinical breast cancer and normal samples from Korean breast cancer patients. In addition, we analysed the mutations, CNAs and gene expression results of approximately 3000 clinical breast cancer samples in the TCGA and METABRIC databases. Based on these study results, we chose the genes that showed a high frequency and recurrent mutations, genomic copy number amplifications and deletions, and expression changes in breast cancer samples, which could be oncogenes, tumour suppressor genes, or breast cancer-associated genes. Among the 121 genes that we chose based on such criteria were previously known oncogenes, tumour suppressor genes and breast cancer-associated genes, including genes involved in DNA repair pathways.

However, our SNUH BCC Panel is unique compared with other cancer panels based on NGS because it includes a certain portion of novel breast cancer-associated genes that have not been included in other recent popular and conventional cancer panels. In this regard, our SNUH BCC Panel is not only targeted to worldwide breast cancer patients but is also ethnically directed to Korean breast cancer patients for diagnosis and therapeutic prognostic prediction.

### Sequence alignment and preprocessing

The NGS sequence reads were mapped to the GRCh37 human reference genome using BWA-MEM^38^ (version 0.7.8) with default parameters. The resulting SAM files are sorted by chromosome coordinates, followed by PCR duplication marking using Picard (version 1.115) (http://picard.sourceforge.net/). Reads with a mapping quality score less than 30 or that have a supplementary alignment were removed from the BAM file before the subsequent analysis.

### Detecting copy number alterations

We used low-depth whole-genome sequencing data and the variable-size binning method^39^ to estimate the CNAs of the samples. Briefly, the whole genome was divided into 10,000 variable-sized bins (median genomic length of bin = 276 kbp) in the case of breast cancer tissue samples and 7,000 bins (median = 396 kbp) in the case of cell line samples, in which each bin had an equal expected number of uniquely mapped reads. Then, each NGS sequence read was assigned to each bin followed by Lowess GC normalization to obtain the read depth of each bin. The copy number was estimated by normalizing the read depth of each bin by the median read depth of the reference DNA.

### Clustering samples based on the CNA dataset

After the CNAs detection followed by MergeLevels^40^, the data underwent multi-sample segmentation with gamma=20. Given the multi-sample segmentation, the event vector was constructed for each cell by providing a value of 0 for the segments with a median copy number of reference DNA, 1 for segments with a gain, and −1 for segments with a loss. Finally, a correlation matrix was constructed, and hierarchical clustering was performed. To evaluate the accuracy of the clustering, a multiscale bootstrapping resampling method was used to approximately calculate the unbiased p-value^41^. The approximately unbiased (AU) p-value indicated the strength of the cluster supported by bootstrapping. Bootstrap samples were generated, and clustering analysis was applied to them repeatedly. The AU p-value of a cluster was related to the frequency that appeared in the bootstrap replicates, and the method was implemented by the R package Pvclust^42^.

### Single Nucleotide Variants Detection

Before SNVs detection, GATK (v3.5-0) IndelRealigner and BaseRecalibrator were used to locally realign reads around the Indel and recalibrate the base quality score of BAM files^43^. We then used three different variant callers (GATK UnifedGenotyper, Varscan, and MuTect) and combined the results to avoid false-positive variant detection^44^. First, GATK UnifiedGenotyper was used with default parameters followed by GATK VariantRecalibrator to obtain filtered variants^43^. Tumour bulk sample and PHLI-seq sorted sample data were processed together to produce a single vcf file. The training data used for variant recalibration included dbSNP build 137, hapmap 3.3, Omni 2.5, and 1000G phase1, and QD, MQ, FS, ReadPosRankSum, and MQRankSum annotations were used for the training. Variants detected in the paired blood sample of the cancer patient were removed to produce the final list of GATK called variants. Varscan2^45^ (ver 2.3.7) and Mutect^46^ (ver 1.1.4) were used with default parameters to produce the lists of Varscan and MuTect called variants, respectively. Here, paired blood read data was also used to separately call germline mutations.

Among the variants detected in the samples by the three variant callers, variants called by at least two callers were collected to obtain intra-sample double called sites. By considering only these variants for subsequent analysis, we could reduce false-positive variant detection derived from NGS errors^44^. Among the intra-sample double called sites, variants found in at least two samples were collected to remove WGA (whole genome amplification) errors, and the genomic loci with the resultant variants were considered confident sites. Finally, a variant in the confident sites was considered to be true if one of the three variant callers detected the variant at the locus and the allele count of the variant was significantly larger than that of the other non-reference bases (Fisher’s exact test, p < 10e-4).

### Single-molecule deep sequencing to validate detected SNVs

We carried out single molecule-deep sequencing^47^ to validate the SNVs detected from the tumour bulk genomic DNA extract. We randomly selected a portion of mutations detected by PHLI-seq and generated a sequencing library for the targeted sites by tagging unique molecular barcodes to each DNA molecule. The difference from the referred paper is that we generated a targeted library using PCR with minimal cycles before tagging the molecular barcodes. This would lose the ability to call duplex consensus sequences (DCSs), but it is still possible to call single-strand consensus sequences (SSCSs). We performed 3-plex PCR using 50 ng of genomic DNA with a 2X KAPA2G Fast Multiplex kit (KAPA Biosystems, KK5802). The amplified products underwent gel electrophoresis, gel purification, and quantification by a Qubit high-sensitivity dsDNA quantification kit (Invitrogen), and normalization for pooling. The pooled library was end-repaired and dA-tailed (NEB, NEBNext^®^ End Repair Module, E6050S). Then, the duplex tags were ligated, and DNA molecules with a length of 200 to 1000 bp were purified to remove dimers. Finally, the molar concentration of the tag-ligated libraries was quantified (**Supplementary Fig. 10**). The constructed library was sequenced by MiSeq 250 PE (Illumina) to generate a 18,182X (median) single-molecule sequencing depth. After generating the SSCSs, the targeted loci (where SNV had been detected) and background loci (where no SNV had been detected) were split. Given the allele frequency distribution from the background group, we prepared the true positive call set from the targeted loci (Benjamini–Hochberg false discovery rate < 0.05).

### Defining cancer genes

After the CNA and SNV detection, potential driver alterations were annotated for cancer-related genes. The gene list consists of 682 genes, which were compiled from The Cancer Gene Census^48^ and The Cancer Gene Atlas (TCGA) Project^27,32^.

